# Multi-organ transcriptomic atlas reveals hallmarks of labor

**DOI:** 10.1101/2024.09.22.614290

**Authors:** Duan Ni, Ralph Nanan

## Abstract

Pregnancy and labor are dynamic processes involving various rapid biological changes. Previous studies found that during pregnancy there are gradual alterations in immune and metabolic pathways. In contrast, changes linked to labor are less well-known, generally limited to reports on specific differentially expressed genes, lacking a more comprehensive overview of changes of the overall signalling landscapes. Moreover, most previous works were standalone, based on individual organs. Here, we aimed to present a more systematic and holistic overview of the impacts of labor on both the maternal and fetal aspects. For this purpose, we collated a multi-organ transcriptomic atlas to decipher the influences of labor on multiple maternal and fetal compartments, covering the maternal adipose tissues (subcutaneous and visceral fat), peripheral blood, myometrium and placenta and fetal cord blood mononuclear cells (CBMCs), and spanning both physiological and pathological conditions. During labor, activation of pro-inflammatory TNF signaling was found in all maternal compartments except adipose tissues, which, along with myometrium, showed upregulation in glycolysis. Myc signaling, as a critical nexus within the intracellular signaling network, was generally enhanced in labor. Contrarily, fetal CBMCs in labor upregulated the anti-inflammatory TGFβ pathway. Importantly, changes in the maternal TNF signaling were conserved across the spectrum of healthy pregnancy of different parities, and pathological conditions like preterm pregnancy and smoking-affected pregnancy. Mechanistically, it might be explained by the reduction in progestin hormonal signals during labor but not by the mechanical stress.

Collectively, we present a comprehensive multi-organ transcriptomic atlas elucidating the maternal and fetal immune and metabolic landscapes associated with labor, shedding light on the complex physiology.

---

Labor signifies the concluding phase of pregnancy, characterised as a dynamic process involving various biological changes influencing multiple organ systems in both the mother and the fetus. While it has been reported that throughout pregnancy there is pronounced maternal immune and metabolic reprogramming^1^, specific changes associated with labor are less studied. In the context of labor, prior studies generally concentrated on individual organ systems^2^, limited to gene-level analyses, attempting to identify specific marker genes^3^. However, comprehensive overviews of signaling pathway changes across different organ systems, spanning both maternal and fetal compartments, are still lacking and needed.

We surveyed on Gene Expression Omnibus for all available transcriptomic datasets across both maternal and fetal compartments to collate a multi-organ transcriptomic atlas cross-sectionally comparing labor versus non-labor (Supplementary Information). The atlas contains 16 datasets, spanning 6 organ systems (maternal blood, maternal subcutaneous fat, maternal visceral fat, placenta, myometrium and cord blood mononuclear cells (CBMCs)), with a total of 392 samples (Figure 1A). Extensive analyses were carried out utilizing these data, focusing on pathway level changes during labor.

**Figure 1.**
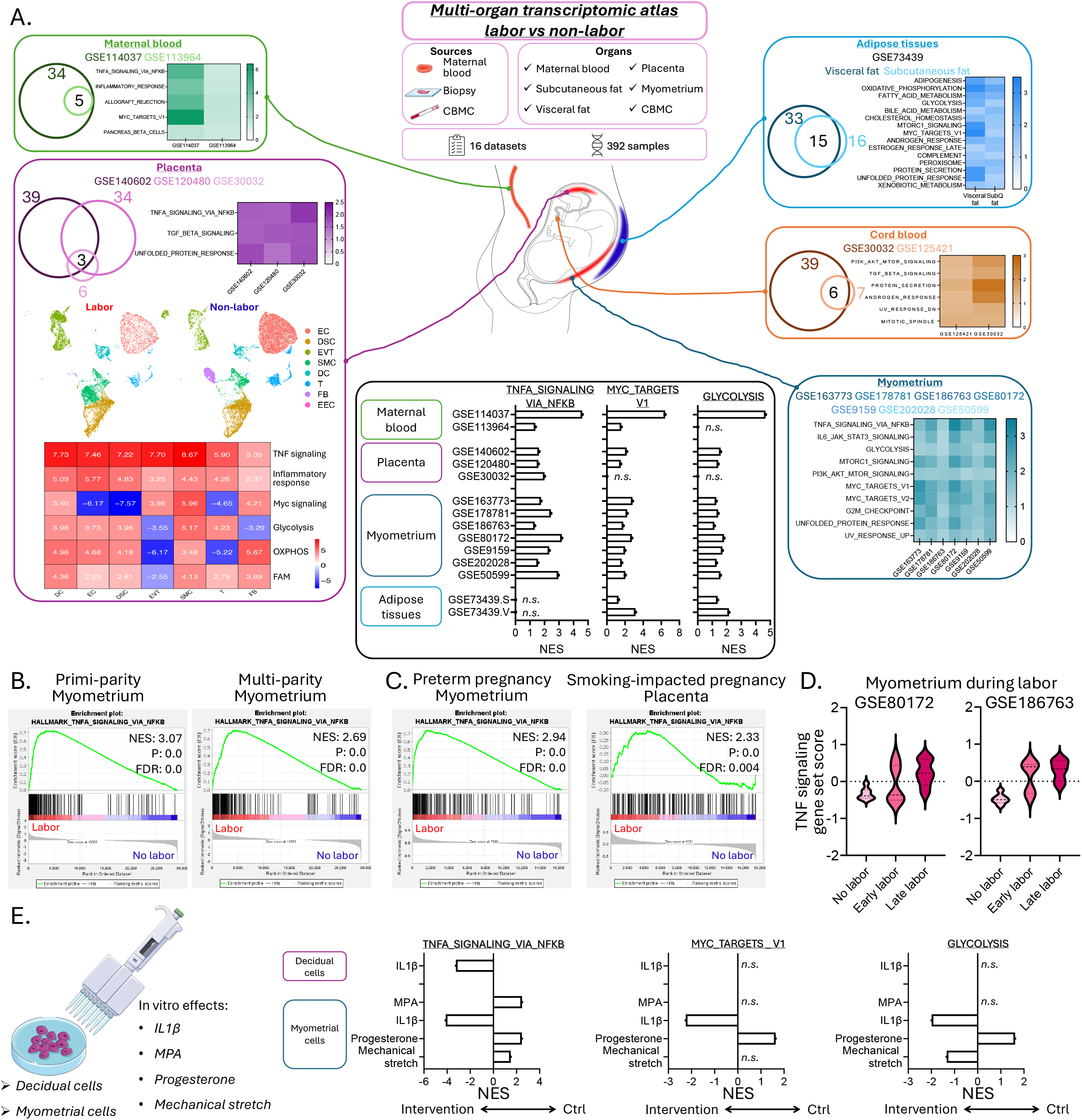
Multi-organ transcriptomic atlas for labor versus non-labor. **A**. A multi-organ transcriptomic atlas was curated from 16 independent datasets comprising of 392 samples, comparing labor versus non-labor through six organ systems (maternal blood, maternal subcutaneous fat, maternal visceral fat, placenta, myometrium and fetal cord blood mononuclear cell (CBMC)). For each organ system, gene set enrichment analysis (GSEA) was run to compare labor versus non-labor. Pathways consistently enriched in labor conditions were depicted by Venn diagrams and heatmaps, with the color reflecting the normalized enrichment score for each condition. In the anatomical graphics, organ systems with prominent changes in immune-related pathways were highlighted in red, the ones with pronounced changes in metabolism-related pathways were highlighted in blue, and the one with mild changes was shown in grey. Representative pathways (TNFA_SIGNALING_VIA_NFKB, MYC_TARGETS_V1 and GLYCOLYSIS) changes across organ systems were summarized in the black box. **B**. GSEA showing the enrichment of TNFA_SIGNALING_VIA_NFKB pathway in primi– and multi-parity myometrium. **C**. GSEA showing the enrichment of TNFA_SIGNALING_VIA_NFKB pathway in preterm pregnancy myometrium and smoking-impacted pregnancy placenta. **D**. Gene set scores of TNFA_SIGNALING_VIA_NFKB pathway in myometrium throughout the labor time course. **E**. Summary of GSEA for representative pathways (TNFA_SIGNALING_VIA_NFKB, MYC_TARGETS_V1 and GLYCOLYSIS) in decidual cells treated with IL-1β and myometrial cells treated with IL-1β, medroxyprogesterone acetate (MPA) and progesterone, and undergoing mechanical stretching *in vitro*.

Various pathway analyses like gene set enrichment analysis (GSEA) were run. For each organ system, we compared the results derived from different datasets and collated the most consistent changes. In maternal blood, labor was linked to upregulation of allograft rejection, TNF-NFκB-related, and Myc-related signaling (Figure 1A). Myc signals were also enhanced in the maternal adipose tissues in labor. This was accompanied with pronounced metabolic changes like enhanced glycolysis, oxidative phosphorylation (OXPHOS) and fatty acid metabolism (FAM) in both visceral and subcutaneous fat (Figure 1A).

We next probed the organs directly implicated in labor like myometrium and placenta (Figure 1A). Similar to adipose tissues, myometrium, which is mainly composed of muscle tissue, also exhibited increased glycolysis and Myc signaling. TNF and IL-6 signaling was higher, possibly induced by mTORC1 activation. These were consistently observed across 7 myometrial datasets.

Immune activation was also found in placenta, as TNF signaling was higher in labor in all 3 datasets (Figure 1A), consistent with previous report^2^. Importantly, a published single cell RNA-seq (scRNA-seq) dataset for placental tissues with/without labor (GSE186368) was re-analyzed (Figure 1A). As in the original study, 8 different cell subsets were identified (endothelial cells, EC; decidual stromal cells, DSC; extravillous trophoblasts, EVT; smooth muscle cells, SMC; dendritic cells, DC; T cells, T; fibroblasts, FB; and endometrial cells, EEC). Endometrial cells were excluded from downstream analysis due to low cellularity. GSEA comparing labor and non-labor found that all cell subsets consistently upregulated the TNF signaling pathway. They also generally displayed more active metabolic profiles including glycolysis, OXPHOS and FAM. An exception were EVTs, where aforementioned metabolic signals were downregulated in labor. Furthermore, Myc signaling was increased in labor in DC, EVT, SMC and FB, but not in EC, DSC and T cells.

Finally, on the fetal side, CBMCs were analyzed (Figure 1A). Intriguingly, the inflammatory and metabolic pathways were generally not affected in CBMCs by labor. Instead, activation of pathways like PI3K-Akt-mTOR pathway and TGFβ signaling was found, the latter usually being linked to anti-inflammatory processes.

Overall, our comprehensive multi-organ transcriptomic atlas analyses revealed that labor significantly shifted the maternal and fetal immune and metabolic landscapes. During labor, activation of pro-inflammatory TNF signaling was found in all maternal compartments except adipose tissues, which, along with myometrium, showed upregulation in glycolysis. Myc signaling, as a critical nexus within the intracellular signaling network, was generally enhanced in labor. Contrarily, fetal CBMCs in labor upregulated the anti-inflammatory TGFβ signaling.

We next interrogated the effects of parity, which is known to influence both the mother and the fetus^4^. As shown in Figure 1B, parity does not confound the impacts of labor, as a consistent increase in TNF signaling was found for myometrium in primi-as well as multi-parity pregnancies. Furthermore, in preterm pregnancies and pregnancies impacted by smoking, labor is also linked to similar patterns of TNF pathway activation (Figure 1C).

Next, using gene set variation analysis (GSVA), we longitudinally analysed two myometrial datasets from early to late labor. They consistently exhibited a gradual increase in TNF signaling throughout the labor time course (Figure 1D).

Finally, we focused on specific perturbations associated with labor, including changes in cytokines, hormones and mechanical stress, in an attempt to decipher the potential underlying causes for aforementioned changes. For this purpose, we curated existing datasets analyzing *in vitro* experiments with decidual or myometrial cells (Figure 1E). Here, IL-1β expectedly activated the TNF signaling in both decidual and myometrial cells but only promoted the Myc and glycolysis pathways in myometrial cells. On the other hand, exposure to progestin hormones, such as medroxyprogesterone acetate (MPA) and progesterone, dampened the TNF signals in myometrial cells. Progesterone also blunted the Myc and glycolysis pathways. In contrast, mechanical stretching of myometrial cells did not increase TNF signaling but promoted glycolysis pathways.

Collectively, these data suggest that the lack of progestin signals, as reflected by their “functional withdrawal” during labor^5^, might be linked to the described transcriptomic changes, whilst mechanical stress on myometrial tissues imposed limited effect.

Here, we present to our knowledge, the first multi-organ transcriptomic atlas comparing labor and non-labor. In-depth analyses with this atlas found that labor was linked to increased TNF-mediated inflammatory signaling in most maternal compartments except adipose tissues but associated with enhanced anti-inflammatory TGFβ signaling in CBMCs. Changes in the maternal compartments seemed to be conserved and were found across various physiological and pathological conditions.

The overall labor-associated increase in TNF signaling was surprisingly not systemic, as it was not found in adipose tissues. However, both adipose tissues and myometrium exhibited enhanced glycolysis, plausibly to supply energy for the labor process. Increased glycolysis might also fuel immune activation^6^ and partly contribute to the increased TNF signaling in maternal tissues.

Mechanistically, these maternal immunometabolic changes could in part be explained by the fall of progesterone signals during labor, while direct cellular mechanical stress surprisingly did not play a role. Other perturbations that might shape the described immunometabolic profiles include changes in hormones like oxytocin and indoleamine 2,3-dioxygenase levels^7^, which warrants further investigation.

The elevated maternal inflammatory signals coincide with the enhanced maternal leukocytosis upon labor^8^. Such immune activation might have evolved to promote maternal immunity against ascending infections during labor and the post-partum period, which remains a major risk accompanying delivery.

Contrarily, the fetal compartment surprisingly exhibited an anti-inflammatory phenotype linked to labor, which might confer protective effects. Inflammation in newborns interferes with multiple physiological processes that are critical for their transition from *in utero* to *ex utero* life^9^. The observed anti-inflammatory TGFβ signaling might thus confer benefits in labor, which might be linked to the protection against pathologies like respiratory distress syndrome and birth asphyxia^10^ in vaginal delivery.

Together, our multi-organ atlas analyses provide novel insights towards the impacts from labor on both the mother and the fetus, shedding more lights on the hallmarks of this sophisticated physiological process.

## Acknowledgements

This project is supported by the Norman Ernest Bequest Fund.

## Author contributions

*Concept and design:* Duan Ni, and Ralph Nanan

*Acquisition, analysis and interpretation of data:* Duan Ni, and Ralph Nanan

*Drafting of the manuscript:* Duan Ni, and Ralph Nanan

*Critical revision of the manuscript for important intellectual content:* All authors

All authors have read and approved the manuscript.

## Conflict of interest

The authors declare no competing interests.

## Notes

### Competing Interest Statement

The authors have declared no competing interest.

