## Supplementary Information for "Multi-organ transcriptomic atlas reveals hallmarks of labor"

### Materials and methods

#### Data curation

We extensively surveyed the Gene Expression Omnibus (GEO) database for studies analyzing labor versus non-labor samples. Notably, studies comparing vaginal delivery and elective caesarean section (no labor) were also included. Datasets included in the current study were listed in Supplementary Table S1.

Single cell RNA-seq (scRNA-seq) data for decidual tissue was downloaded from GEO database (GEO ID: GSE186368)[1] and analyzed with *Seurat* (v4)[2].

**Table S1.** Transcriptomic datasets used in the current study.

| **Physiological conditions** | | | |
|---|---|---|---|
| **Organ system** | **Sample #** | **GEO ID** | **Ref** |
| Myometrium | 5 labor vs 5 non-labor | GSE163773 | [3] |
| | 8 labor vs 8 non-labor | GSE178781 | [4] |
| | 7 labor vs 5 non-labor | GSE80172 | [5,6] |
| | 3 labor vs 3 non-labor | GSE9159 | [7] |
| | 7 labor vs 5 non-labor | GSE186763 | [8] |
| | 4 labor vs 7 non-labor | GSE202028 | [9] |
| | 5 labor vs 5 non-labor | GSE50599 | [10] |
| Subcutaneous fat | 15 labor vs 25 non-labor | GSE73439 | [11] |
| Visceral fat | 15 labor vs 25 non-labor | GSE73439 | [11] |
| Maternal blood | 8 labor vs 8 non-labor | GSE113964 | [12,13] |
| | 8 labor vs 8 non-labor | GSE114037 | [12,13] |
| Placenta | 31 vaginal delivery vs 3 c-section | GSE30032 | [14] |
| | 6 spontaneous labor vs 6 elective c-section | GSE120480 | [15] |
| | 16 labor vs 18 non-labor | GSE140602 | - |
| CBMC | 8 vaginal delivery vs 7 c-section | GSE125421 | [16] |
| | 22 vaginal delivery vs 2 c-section | GSE30032 | [14] |

| **Pathological conditions** | | | | |
|---|---|---|---|---|
| **Organ system** | **Sample #** | **Disease conditions** | **GEO ID** | **Ref** |
| Myometrium | 8 labor vs 8 non-labor | Preterm birth | GSE134447 | [17] |
| Placenta | 16 vaginal delivery vs 7 c-section | Pregnancy exposed to smoking | GSE30032 | [14] |

| *In vitro* experiments | | | |
|---|---|---|---|
| **Cell type** | **Sample #** | **GEO ID** | **Ref** |
| Primary myometrial cells | 6 ctrl vs 3 progesterone vs 3 IL-1β | GSE68171 | 18 |
| Primary myometrial cells | 9 ctrl vs 9 medroxyprogesterone acetate | GSE22961 | 19 |
| Decidualized human endometrial stromal cells | 3 ctrl vs 3 IL-1β | GSE58220 | 20 |

**Data analysis**

Gene set enrichment analysis (GSEA) was run with the software (v4.3.2) following the developer's tutorial[21]. Gene set score was calculated using gsva (v1.51.5) package[22] based on the "Human gene set: HALLMARK_TNFA_SIGNALING_VIA_NFKB" gene set from GSEA software. For scRNA-seq analysis, *Seurat* package was used[2]. In brief, data was first quality checked and then integrated with *LIGER*[23] to minimize potential batch effects. Cells were clustered using marker genes reported in the original publication[1]. GSEA was run with the *fgsea* package.
